# Detection of Reticuloendotheliosis Virus in Brazil

**DOI:** 10.1101/698985

**Authors:** Giovana S. Caleiro, Cristina F. Nunes, Paulo R. Urbano, Karin Kirchgatter, Jansen de Araujo, Edison Luiz Durigon, Luciano M. Thomazelli, Brittany M. Stewart, Dustin C. Edwards, Camila M. Romano

## Abstract

Reticuloendotheliosis retroviruses (REV) are known to cause immunosuppressive and oncogenic disease that affects numerous avian species. REV is present worldwide and recently has been reported in South America with cases of infected commercial flocks in Argentina. We surveyed for the presence of REV in birds from a state in the northern region of Brazil using real-time PCR. We report the first cases of REV in Brazil, detected in Muscovy ducks (*Cairina moschata*), wild turkeys (*Meleagris gallopavo*), and chickens (*Gallus gallus*) at a relatively high prevalence rate (16,8%). Phylogenetic analysis indicated a close relationship of this strain to variants in the United States. This study provides evidence of REV in the Amazon biome and provides a baseline for future surveillance of the virus in the region and throughout Brazil.

## Main Text

Reticuloendotheliosis viruses (REV) are a group of immunosuppressive and sometimes oncogenic retroviruses that affect numerous species of birds, including waterbirds (Anseriformes), game birds (Galliformes), and perching birds (Passeriformes) (Nair et al. 2013). REV has previously affected commercial poultry and has been a recurring obstacle in the conservation of endangered species (Luan et al. 2016). Within vaccinated flocks, REV infection can lead to decreased efficacy of vaccinations for avian influenza virus, Newcastle disease virus, Marek’s Disease virus (MDV), and turkey herpesvirus due to the reduced humoral response resulting from immunosuppression (Sun et al. 2009). The virus was first identified in the US in 1958, and later in China, Taiwan, Australia, Argentina, and Canada (Singh et al. 2003; MacDonald et al. 2018). REV prevalence varies from 0–50% depending on the region, setting, and avian species (Jiang et al. 2013; Ferro et al. 2017).

Evidence of REV in South America was recently demonstrated in fowlpox-vaccinated commercial poultry flocks in Argentina (Buscaglia, 2013). REV proviral DNA has been detected in the genomes of the attenuated MDV vaccine (MD-2 strain) and the field and vaccine strains of fowlpox viruses (FWPV-REV) (Isfort et al. 1992; Hertig et al. 1997). Vaccine-integrated REV can be either infectious or non-infectious (Hertig et al. 1997; Moore et al. 2000). The Brazilian coast serves as an important stopover site for migratory birds coming from the northern hemisphere. Several species migrate during the austral winter from Argentina and Chile to Central Bolívia and Brazil (Somenzari et al. 2018). In addition, several birds, including passeriformes, migrate from the US, Mexico, and Central America to northern Brazil (Somenzari et al. 2018). However, no prior studies have surveyed for the presence of REV in Brazil. With more than 2,000 species of wild birds present in the country, many of which are endangered, determining the presence of REV and establishing a baseline prevalence rate could be of value for future conservation efforts in the Amazon biome.

During 2005–2006, a total of 441 samples were collected (blood and pooled cloacal/orotrachea swabs) near eight different cities in the northeastern region of Pará state. Most of the samples were from Muscovy ducks (*Cairina moschata*, *n*=379), and the remaining samples were from turkeys (*Meleagris gallopavo*, *n*=41), and chickens (*Gallus gallus*, *n*=21). Total cDNA from orotrachea and cloacal swab samples was obtained and previously prepared from an avian influenza virus detection study (Thomazelli et al. 2012). As an internal control, a conventional PCR method for avian mitochondrial DNA was performed using cytochrome *b* primers (Kocher et al. 1989). The presence of REV proviral cDNA was detected using real-time PCR and primers specific to the *gp90* gene (*env*) (Li et al. 2013). Reactions were prepared using 2.5mM of each primer, 7.5μL of water, 12.5μL of MasterMix SYBR® Green (LifeTechnologies^®^, Brazil), and 3μL of purified cDNA. Plasmids containing genes of interest were constructed from commercially-synthesized inserts (GenScript, USA) to serve as positive controls. In addition, partial fragments of LTR-U5, Gag, and envelope from REV-positive samples were amplified and sequenced by Sanger method using previously described primers (Singh et al. 2003; Barbosa et al. 2007; Li et al. 2013). For the envelope region, an additional primer was designed (gp90-7242R 5’–GCCAGTATGCACAGCCCTATCCA–3’).

Of the 441 samples tested and amplified, REV PCR products were detected in 74 samples (16,7%). Infected individuals included 65 Muscovy ducks, 6 wild turkeys, and 2 chickens. REV positive samples came from Marabitana, Vila Maracajá, and Marajó. Sequencing reads were inspected for quality and consensus sequences were built using CLC genomic workbench v5 (https://www.qiagenbioinformatics.com/) and submitted to GenBank, accession numbers MG953804–MG953809. Consensus sequences of partial LTR/gag (808 bp), gag (1168 bp), and envelope (~600 bp) were aligned and compared to reference strains and other sequences retrieved from GenBank representing worldwide REV (*n*=16). Phylogenetic trees were reconstructed under maximum likelihood method in PhyML software using K80 as the best nucleotide substitution model for all fragments as determined by jModeltest (Guindon et al. 2010; Darriba et al. 2012). Only sequences built with high quality reads (phred >30) were used for phylogenetic reconstructions. Genetic distance (*p*-distance) between the main REV strains and one representative Brazilian REV sample was estimated for LTR/gag, gag, and envelope partial fragments. To establish the phylogenetic relationships among Brazilian and worldwide REV, phylogenetic trees were reconstructed using LTR/gag and envelope fragments representing Brazilian and worldwide REV genomic sequences (*n*=16) (Fig 1). Brazilian REV clustered together with sequences sampled in the US, such as APC-566. Genetic distance at the nucleotide level (*p*-distance) between the main REV strains and one representative Brazilian REV also agreed that Brazilian REV is more closely related to the US samples representative of subtype 3, such as MD-2 and APC-566, rather than to spleen necrosis virus (SNV) strain or China isolates (Fig. 1 and Table 1).

**Figure 1.**
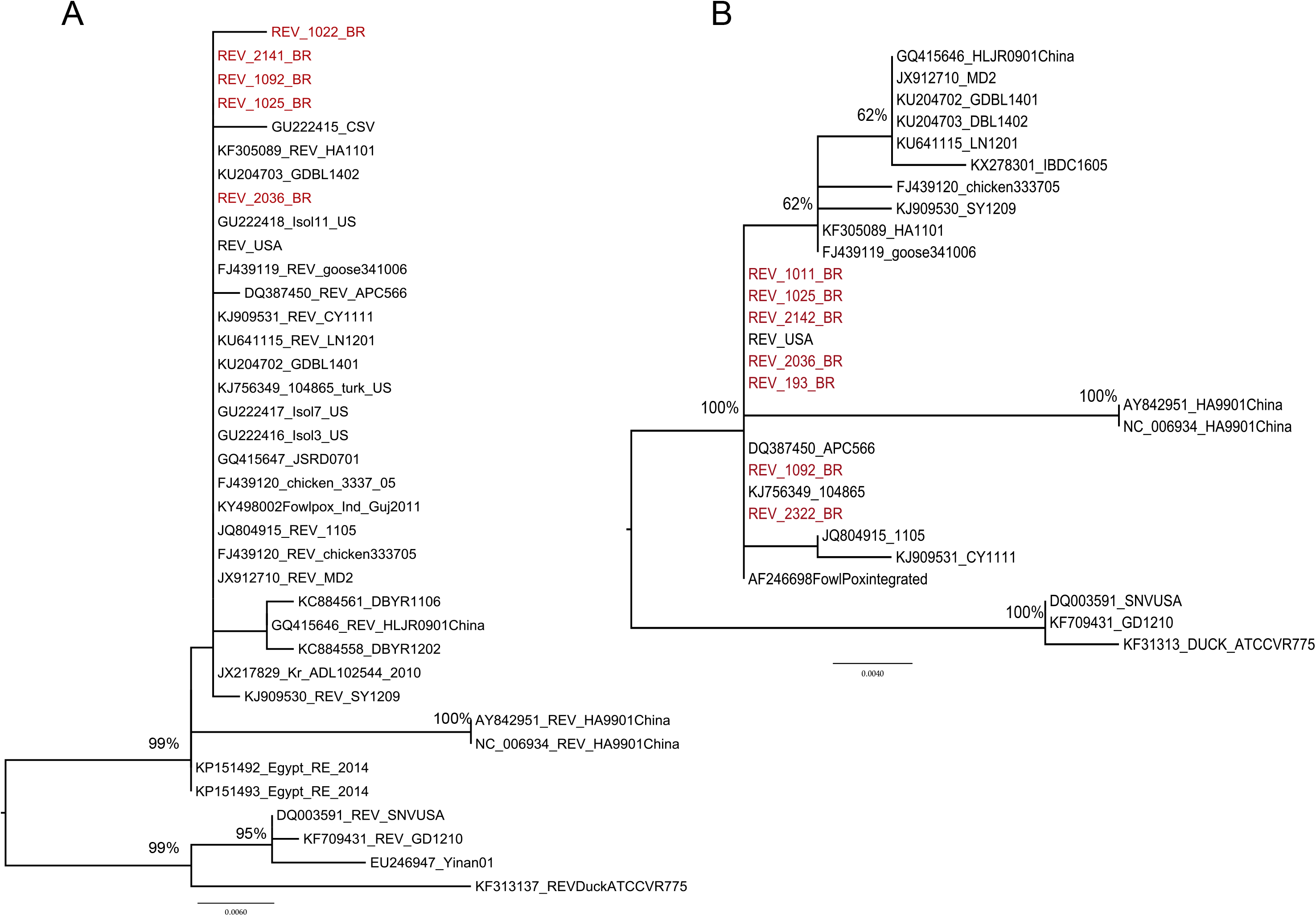
Maximum likelihood Phylogenetic reconstruction showing the relationships between REV from Brazil and other isolates. Brazilian sequences (in red) are from Muscovy ducks (*Cairina moschata*). The trees were made with partial sequences from REV env (A) and LTR/gag (B) available at GenBank.

**Table 1.** Genetic Identity at nucleotide level between Brazilian REV and reference strains.

| REV_2036_PA_Br |  |  |  |  |
| --- | --- | --- | --- | --- |
| Reference | GenBank ID | LTR/gag | Gag | Env |
| APC-566 | DQ387450 | 99.8% | 100% | 99.8% |
| SNV | DQ003591 | 96% | 97.5% | 96.4% |
| MD-2 | JX912710 | 99.4% | 100% | 100% |
| FPV | AF246698 | 99.6% | 100% | 100% |
| CSV | DQ237905 | NA<sup>a</sup> | NA | 99.5% |
| HA9901 (China) | AY842951 | 96% | 98% | 97.6% |

Distance matrix (*p*-distance)
<sup>a</sup>NA – sequence not available

Pará state is located in the northern coastal region of Brazil and has a tropical rainforest climate. Due to its geographic location and ecosystem, Pará is a stopover site for shorebirds coming from the northern hemisphere during the migration season. In addition, several species of migratory birds that have Argentina and the US as their final destination pass through Brazil during their flight. The similarity between Brazilian REV and the US viruses suggests that the US could be the source of REV detected in Brazil. However, there is no sequence data available for viruses found in Argentina. As such, the source of Brazilian REV cannot be definitively determined. Representative strains of REV include the defective REV-T, the non-defective REV-A, SNV, duck infectious anemia, and chicken syncytial virus (CSV), and the MD-2 and FWPV vaccine-integrated strains. As there is relatively little genetic variation among REV strains, it is difficult to determine the origin and classification of an isolate (Bohls et al. 2006). In our analysis, Brazilian REV differs only 0.0–0.5% from both FWPV and MD-2-integrated REVs and non-integrated APC-566 and CSV. We inspected for point mutations, which have been characterized as specific for FWPV-REV (Tadese et al. 2008). According to the genetic pattern observed, the Brazilian REV samples are all non-FWPV-integrated strains. However, not all FWPV-REV envelope sequences available on GenBank (KY498002, AF246698, and JX217830) contain this signature, indicating that some genetic variability among integrated strains exists and determining the integration status based on few point mutations is inaccurate (Tadese et al. 2008). Therefore, it remains undetermined if Brazilian REV is present as infectious particles or is instead integrated within a large DNA virus genome.

Reticuloendotheliosis viruses can infect a number of species, including captive and wild perching birds, game birds, and waterbirds. Although we detected REV in Muscovy ducks, wild turkeys, and chickens, we could only amplify and sequence the virus from ducks. It is possible that the virus was present in a low viral load in other species, and we could not generate large amplicons. Muscovy ducks are native to Mexico, Central, and South America, but populations of Muscovy ducks reside in the US, mainly in Florida and southern Texas. These birds are essentially non-migratory or irregular migrants without any established migration patterns, only migrating short distances to avoid dry weather and fluctuating water conditions. Chickens and turkeys, frequently associated with REV infections, are also non-migratory birds. Therefore, we consider it is unlikely that the positive birds found in Brazil were infected elsewhere.

Here, we detected REV in Brazil for the first time and presented the first DNA sequences of REV provirus from South America. REV found in Brazil is similar to other common circulating strains, including those found in the US. Additionally, considering that REV was present in samples from three of the five regions collected from Pará from 2005–2006 (located 200–500 km apart from each other), it is very likely that the virus has spread to other states in Brazil. We hypothesize that the virus could be carried by migratory birds which stopover in the northern part of the country on their way to and from North America. These results suggest the need for additional studies to further determine the prevalence and genetic variability of REV in Brazilian wild and captive birds and also to perform risk assessment studies dedicated to free-range and captive commercial avian species.

All procedures involving wild and captive birds were approved by the Animal Ethics Committee from the Instituto de Medicina Tropical de São Paulo under protocol 000283A, and licensed by the Ministério do Meio Ambiente-MMA at the Instituto Chico Mendes de Conservacão da Biodiversidade (ICMBio/SISBIO) under protocols 34605-7 45527-2, 54616-1, and 201/2006 CGFAU. This work was supported by Fundação de Amparo a Pesquisa do Estado de São Paulo (FAPESP Project #2015/05958-3). G. Caleiro holds a CAPES scholarship.

